# Arousal-dependent auditory responses in the brain of Bengalese finches measured by gene expression

**DOI:** 10.1101/2021.06.01.446331

**Authors:** Takafumi Iizuka, Chihiro Mori, Kazuo Okanoya

## Abstract

Songbirds use auditory feedback to maintain their own songs. Juveniles also memorize a tutor song and use memory as a template to make up their own songs through auditory feedback. A recent electrophysiological study revealed that HVC neurons respond to BOS playback only in low arousal, sleeping, or anesthetized conditions. One outstanding question is how does auditory suppression occur in the brain? Here, we determined how arousal affects auditory responses simultaneously in the whole brain and over the song neural circuit in Bengalese finches, using the immediate early gene *egr-1* as a marker of neural activity. Our results showed that auditory responses in the low-arousal state were less susceptible to gating, which was also confirmed by gene expression, and that the suppression may be weaker than observed in previous zebra finch studies. This may be because the Bengalese finch is a domesticated species. In addition, our results suggest that information may flow from the MLd.I of the midbrain to higher auditory regions. Altogether, this study presents a new attempt to explore the auditory suppression network by simultaneously investigating the whole brain using molecular biology methods.

## 1. Introduction

Songbirds use auditory feedback to maintain their songs. Juveniles also memorize a tutor song and use memory as a template to make up their own songs through auditory feedback (Okanoya & Yamaguchi, 1997; Tumer & Brainard, 2007). A recent electrophysiological study revealed that HVC (proper name) neurons only respond to Bird’s own songs (BOS) playback in low arousal, sleeping, or anesthetized conditions. The gating phenomenon, in which neurons respond to BOS playback only during sleep or under anesthesia, is the sole known evidence for controlling auditory input into the song system (Cardin & Schmidt, 2003, 2004; McCasland & Konishi, 1981; T. A. Nick & Konishi, 2001; Teresa A. Nick & Konishi, 2005; Rauske, Shea, & Margoliash, 2003; Schmidt & Konishi, 1998). In the songbird, the auditory input is transmitted from the inner ear to the primary auditory regions, such as MLd (dorsal part of the lateral mesencephalic nucleus) and Ov (nucleus ovoidalis) in the midbrain; to the lower auditory regions, such as field L; higher auditory regions, such as NCM (caudomedial nidopallium) and CMM (caudal medial mesopallium) in the cerebrum; and finally to the HVC. Although the mechanism of auditory suppression remains unclear, it is generally believed that auditory input suppression is necessary to prevent information overload (Cromwell, Mears, Wan, & Boutros, 2008; Freedman et al., 1987). Auditory input suppression in songbirds has received much attention because it may be involved in the control of auditory feedback that is required for vocal learning and song maintenance (Schmidt & Konishi, 1998).

When an action potential is generated in a neuron, voltage-gated calcium channels open, and calcium ions flow into the cell. The influx of calcium ions activates intracellular proteins and other signals and induces gene expression in the nucleus. In particular, the expression of immediate early genes is induced within a very short time after the action potential is generated, and the level of immediate early gene expression reflects the neural activity observed in electrophysiological experiments (Jarvis & Nottebohm, 1997; Mello, Vicario, & Clayton, 1992). Therefore, gene expression has been used as a marker of neural activity. Early growth response protein 1 (*EGR-1*), the first gene evaluated in this study, is expressed in the nucleus 5 min after the onset of an action potential and is transferred to the cytoplasm 30 min later (Velho, Pinaud, Rodrigues, & Mello, 2005). Here, we investigated the relationship between gating and arousal conditions, with a focus on the immediate early gene expression that is dependent on neural activity in the whole brain of songbirds. Our findings verify the relationship between auditory information flow and auditory suppression.

## 2. Materials and Methods

### 2.1 Subjects

Most male Bengalese finches (BFs, Lonchura striata var. domestica) were obtained from a local breeder (n =21), while others were laboratory bred (N = 5). The photoperiod was maintained at a 14:10 h light/dark cycle, with food and water provided ad libitum. The original research reported herein was performed under the guidelines established by the Institutional Animal Care and Use Committee of the University of Tokyo.

### 2.2 Stimuli

BOS were presented to each bird. Before the experiments, bird vocalizations were recorded to generate auditory stimuli. Recordings were conducted in sound-proof chamber (11.0 cm×11.0 cm×12.0 cm) using a microphone (PRO 35, Audio-Technica, China) connected with an audio-interface (Octo-capture Ex, Roland, Japan), with a sampling rate of 44.1 kHz. To create BOS playback, we chose one segment of each bird’s song from the entire series of recorded vocalizations. The segment chosen was approximately 15 s and contained all song notes. Songs were automatically saved using Avisoft SASLab Pro software (Avisoft Bioacoustics, Germany).

### 2.3 Experiment

The arousal level was defined by the experimental treatments. Three conditions: awake condition; immediately after lightning, sleep condition; 2h after turning light off, and anesthetized condition; after anesthetization, were evaluated. Song stimuli (pseudorandom sequence) were presented to birds for 15 s, and the inter-onset interval (IOI) was 1 min. The BOS was played 30 times at about 65 dB from a speaker before ending. Silent controls were also performed under the three conditions. In anesthetized conditions, a 20% urethane solution of about 70 uL was injected intraperitoneally 10 min before the experiment began. We confirmed the bird’s breathing stability and closed eyes at the beginning and end of the experiment.

### 2.4 History

For brain sampling, the birds were euthanized by decapitation. Brains were removed and immediately embedded in OCT compound (Sakura Fine Tech, Tokyo, Japan) inside tissue block mounds, frozen on dry ice, and stored at -80 °C until use. Frozen sections (12-μm thick) were cut in the sagittal plane using a cryostat (Leica, Germany).

### 2.5 In-situ hybridization

The experimental operation was based on the protocol described by Wada *et al*. 2013 *Eur J Neurosci*, using the same *egr-1* RNA probe as used by Hayase *et al*. 2018 *PLOS Biol* (Hayase et al., 2018), which was labeled with digoxigenin (DIG). Brain sections were fixed in 4% paraformaldehyde/ 1×PBS (pH 7.4), washed in 1×PBS, acetylated, dehydrated in an ascending ethanol concentration series, air-dried, and processed for *in situ* hybridization with antisense DIG-labeled *egr-1* riboprobes. A total of 265 ± 44.83 (mean ± SD) ng of the *egr-1* probe was added to a hybridization solution (50% formamide, 10% dextran, 300 mM NaCl, 10 mM Tris-HCl (pH 8.0), 12 mM EDTA (pH8.0), 0.1% N-Lauroylsarcosone, 0.2 mg/mL tRNA, 10 mM dithiothreitol and 1×Denhardt’s solution). Hybridization was performed at 70 °C for 13 h. The slides were washed in 5×SSC at 65 °C for 30 min, 4×SSC and 50% formamide at 65 °C for 40 min, 2×SSC and 50% formamide at 65 °C for 40 min, 0.1×SSC at 65 °C for 15 min twice, 0.1×SSC at room temperature for 15 min, NTE buffer at room temperature for 20 min, and 1×TNT buffer at room temperature for 5 min. To remove endogenous alkaline phosphatase, the slides were washed in 0.6% H2O2 and 1×TNT buffer at room temperature for 30 min. The slides were then washed three times with 1×TNT buffer at room temperature for 5 min. To prevent non-specific binding, the slides were treated with 120 uL of DIG blocking solution at room temperature for 30 min, followed by a wash with 1×TNT buffer for 1 s. Slides were then treated with peroxidase-conjugated anti-DIG antibody/ DIG blocking solution (1:500) at 4 °C for 20 h and then washed three times with 1×TNT buffer, at room temperature for 5 min. The slides were then treated with fluorescein/ 1×plus amplification diluent (1:200) at room temperature for 10 min and washed twice in 1×TNT buffer for 5 min, 1×TNT buffer for 20 min, and 1×TNT buffer for 5 min. The slides were dipped into milliQ water for 1 s and mounted with VECTASHIELD Mounting Medium containing DAPI.

### 2.6 Analysis

To quantify the *egr-1* mRNA signal, brain fluorescence images were taken with a microscope (BZ-X700, KEYENCE, Osaka, Japan) with a 40X/0.95NA objective lens (PlanApoλ, Nikon, Tokyo, Japan), GFP and DAPI filters, and a monochromatic cooled-CCD camera. The images were quantified using the macro-cell-count function of the BZ-X analysis application.

### 2.7 Statistical analysis

A statistical analysis of the effects of auditory stimulation and arousal manipulation on *egr-1* expression was performed using two-way analysis of variance. In addition, a correlation analysis of regional activities was performed using the Spearman’s rank correlation test.

## 3. Results

We found significant treatment effects in certain brain regions. *egr-1* expression significantly increased upon receiving auditory stimuli in HVC (F = 52.04, p = 2.85 ×10^−8^, η^2^= 0.61), NCM (F = 8.27, p= 9.69 ×10^−3^, η^2^= 0.30), MLd.O (F = 5.7, p = 0.024, η2= 0.12), MLd.I (F = 10.79, p = 2.91 ×10^− 3^, η^2^= 0.21), and there was an arousal effect in MLd.O (F = 11.71, p = 0.002, η^2^= 0.25), MLd.I (F = 13.71, p = 1.01 ×10^−3^, η^2^= 0.27).

There were also correlations between HVC and NCM in the awake condition (r = 0.886, p = 0.033), HVC and MLd.O in the sleep condition (r = 0.708, p = 0.050), HVC and MLd.I during sleep (r = 0.781, p = 0.022) and awake (r = 0.886, p = 0.033_ conditions, NCM and X in sleep condition (r = 0.874, p = 0.0045), NCM and MLd.I in sleep condition (r = 0.884, p = 0.0036), MLd.I in sleep condition (r = 0.756, p = 0.030), and MLd.O and MLd.I in sleep (r = 0.775, p = 0.024) and awake (r = 0.886, p = 0.033) conditions. In addition, there was a significant correlation between HVC and NCM in the sleep condition (r = 0.695, p = 0.056), HVC and X in the sleep condition (r = 0.714, p = 0.058), HVC and MLd.O in the awake condition (r = 0.829, p = 0.058), and NCM and MLd.I in the awake condition (r = 0.829, p = 0.058).

## 4. Discussion

There were significant changes in gene expression in response to auditory stimuli, but no difference was observed in response to arousal manipulation in the HVC and NCM. In the HVC, arousal might be absent in the Bengalese finch compared with a sleeping zebra finch. In X, no auditory response was observed, regardless of the condition. The Bengalese finch is a domesticated songbird that may have weaker gating than zebra finches, as suggested by the results of other electrophysiological experiments (Prather et al., 2008). Both the inner and outer sides of the MLd are affected by arousal manipulation, with higher activity during wakefulness than during sleeping. In the lower auditory cortex, the higher activity during wakefulness may be due to the lack of upstream auditory information gating. These results are consistent with recent research on electrophysiological gating techniques in songbirds (Cardin & Schmidt, 2003, 2004; McCasland & Konishi, 1981; T. A. Nick & Konishi, 2001; Teresa A. Nick & Konishi, 2005; Rauske et al., 2003; Schmidt & Konishi, 1998).

There was also a correlation between MLd.O and MLd.I, regardless of arousal manipulation and activities. In contrast, there was no correlation or correlative trend between the HVC and X in the awake condition. This means that auditory information activities are linked in other regions, up to X, which includes the correlative trend in the sleep condition. In addition, in the awake condition, auditory information activities are linked up to the HVC. Thus, gating occurs in the song nucleus under awake conditions. This correlation trend has been confirmed by an increasing the number of individuals.

The inner and outer properties and projection relationships have been determined from sound source localization experiments using barn owls (Takahashi & Konishi, 1988) and are used in different paths. On the other hand, there might not be a strict separation between the inner and outer projection in the zebra finch because they overlapped significantly (Krützfeldt, Logerot, Kubke, & Wild, 2010; Logerot, Krützfeldt, Wild, & Kubke, 2011; Martin Wild, Krützfeldt, & Fabiana Kubke, 2010). In addition, electrophysiological experiments have shown dorsoventral tonotopy, with low frequencies being expressed dorsally and high frequencies ventrally (Woolley & Casseday, 2004). MLd inner and outer differences remain unknown in songbirds, but in this study, auditory responses were higher during wakefulness than during sleeping, and there was a correlation between regions in both conditions, suggesting that auditory responses may interact with MLd. On the other hand, there was no correlation between MLd.O and NCM, regardless of the condition, suggesting that auditory information flows from MLd.I to the higher auditory cortex.

This study suggests that auditory information flows from the lower auditory cortex to the higher differs under arousal, sleep, and awake conditions, based on our gene expression analysis of the whole brain. This method is useful for detecting many neural activities simultaneously. Here, we confirmed that the gating phenomenon could be detected by using neural markers in an electrophysiological and biological approach.

## 6. Figure

**Figure. 1.**
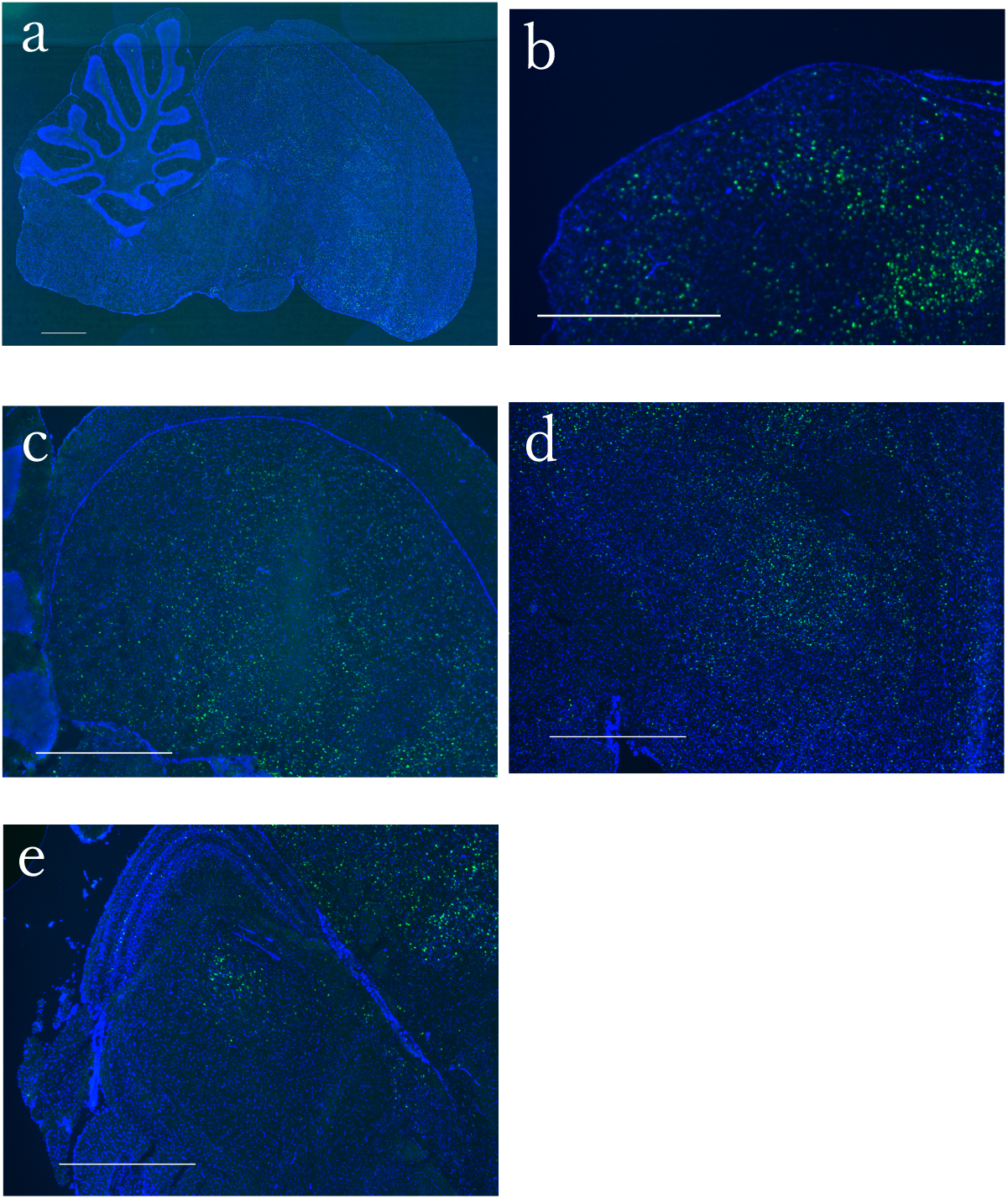
Representative fluorescence *in-situ* hybridization result of *egr-1* expression in the Bengalese finch’s brain. Sagittal sections of the whole brain following *in situ* hybridization are shown. Green signals (fluorescein) indicate *egr-1* mRNA localization and blue signals (DAPI) indicate nuclei. (a) Whole brain, (b) HVC, (c) NCM and other higher auditory cortex, (d) X and the surrounding striatum, (e) MLd and the surrounding midbrain; scare bar = 1 mm.

**Figure. 2.**
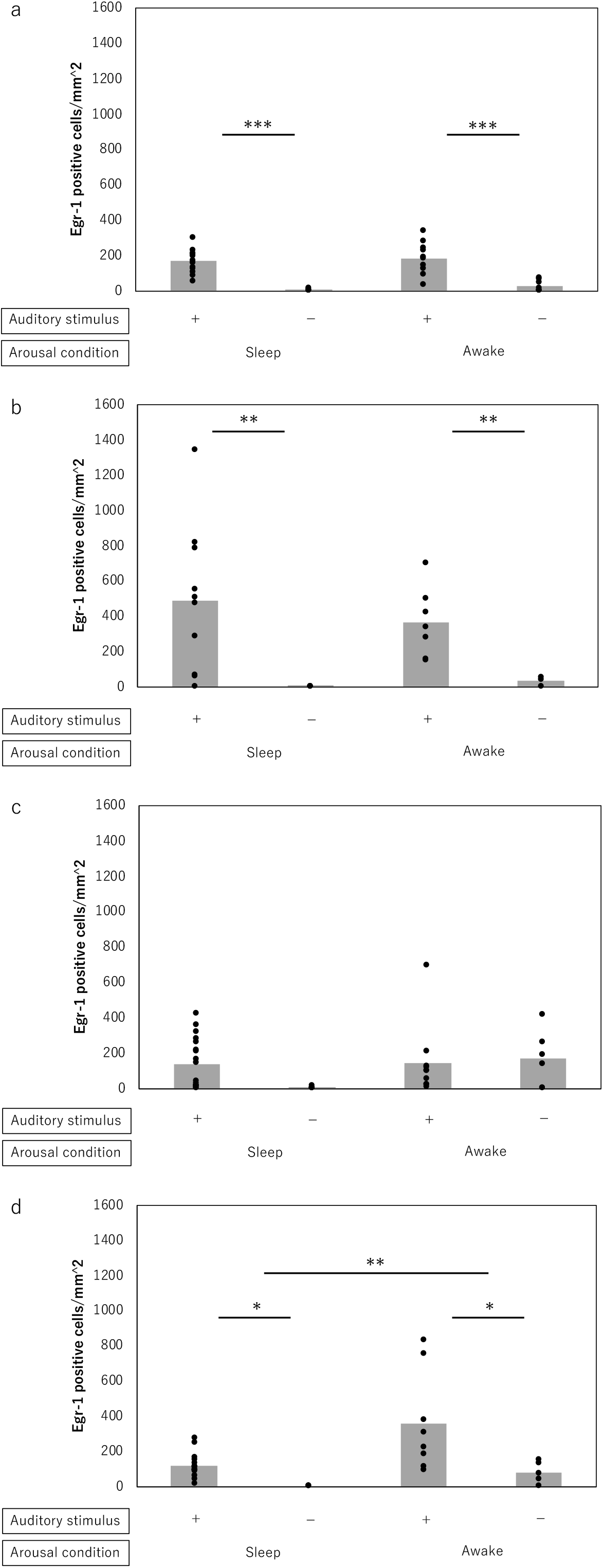
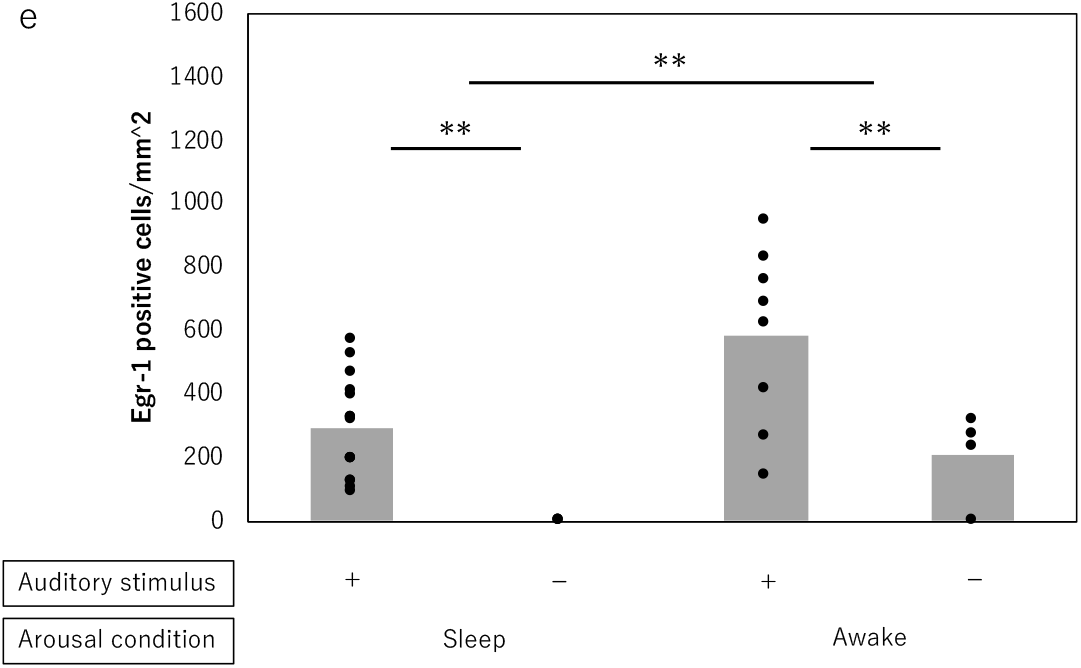
*egr-1* expression following auditory stimuli and arousal conditions. The density of positive cells is presented. Each plot shows the individual data, and each bar represents the average. (a) HVC (properly named), (b) NCM (caudomedial mesopallium), auditory cortex, (c) X, striatum, (d) MLd.O (dorsal part of the lateral mesencephalic nucleus, outer), (e) MLd.I (dorsal part of the lateral mesencephalic nucleus, inner). Data are means; *** p < 0.001, ** p < 0,01, * p < 0.05, ANOVA.

**Figure. 3.**
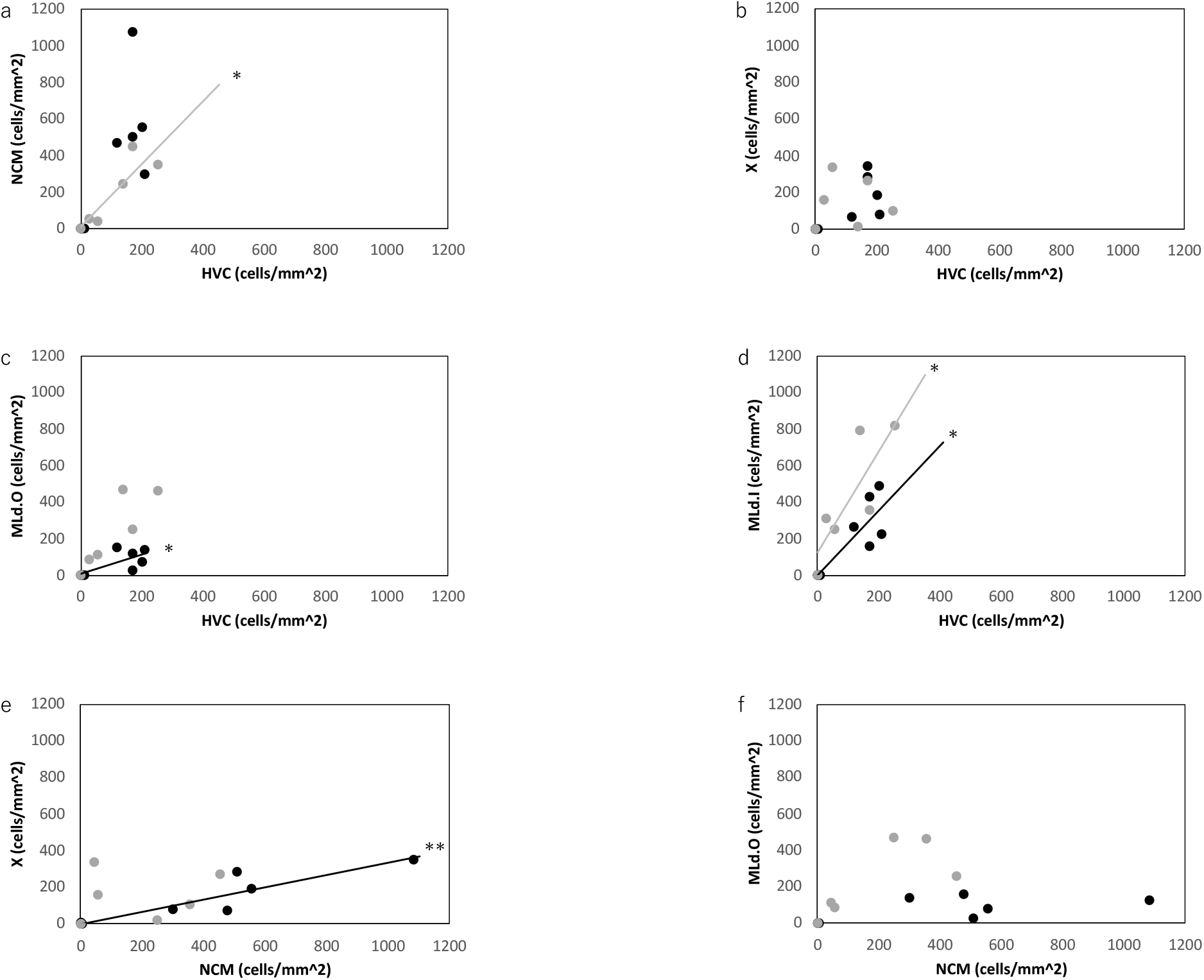
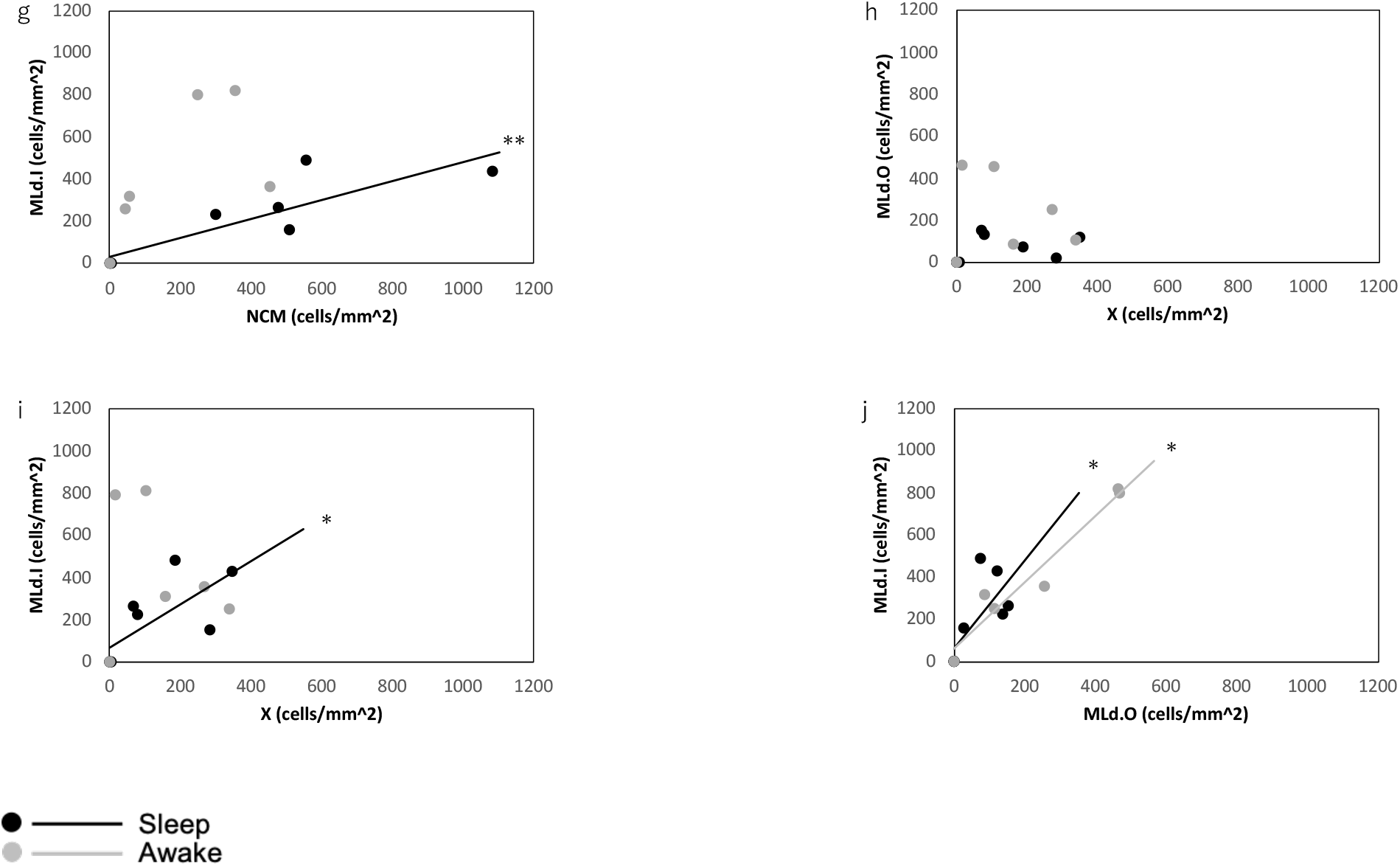
Quantification of *egr-1* expression correlation in two song nuclei (area X and HVC) and two auditory relays (NCM and MLd). A closed circle indicates sleep condition, and a glay circle indicates awake condition. These plots show the correlation between *egr-1* expression in certain brain areas.

